# Continuous and bolus intraventricular topotecan prolong survival in a mouse model of leptomeningeal medulloblastoma

**DOI:** 10.1101/443994

**Authors:** GM Shackleford, MY Mahdi, RA Moats, Debra Hawes, HC Tran, T Hoang, E Meng, A Erdreich-Epstein

## Abstract

Leptomeningeal metastasis remains a difficult clinical challenge. Some success has been achieved by direct administration of therapeutics into the cerebrospinal fluid (CSF) circumventing limitations imposed by the blood brain barrier. Here we investigated continuous infusion versus bolus injection of therapy into the CSF in a preclinical model of human Group 3 medulloblastoma, the molecular subgroup with the highest incidence of leptomeningeal disease. Initial tests of selected Group 3 human medulloblastoma cell lines in culture showed that D283 Med and D425 Med were resistant to cytarabine and methotrexate. D283 Med cells were also resistant to topotecan, whereas 1 μM topotecan killed over 99% of D425 Med cells. We therefore introduced D425 Med cells, modified to express firefly luciferase, into the CSF of immunodeficient mice. Mice were then treated with topotecan or saline in five groups: continuous intraventricular (IVT) topotecan via osmotic pump (5.28 μg/day), daily bolus IVT topotecan injections with a similar daily dose (6 μg/day), systemic intraperitoneal injections of a higher daily dose of topotecan (15 μg/day), daily IVT pumped saline and daily intraperitoneal injections of saline. Bioluminescence analyses revealed that both IVT topotecan treatments effectively slowed leptomeningeal tumor growth in the brains, although histological analysis showed that they were associated with localized brain necrosis. In the spines, bolus IVT topotecan showed a trend towards slower tumor growth compared to continuous (pump) IVT topotecan, as measured by bioluminescence. Both continuous and bolus topotecan IVT showed similar survival that was longer compared to other groups. Thus, both direct IVT topotecan CSF delivery methods produced better anti-medulloblastoma effect compared to systemic therapy at the dosages used here.

## Introduction

Medulloblastomas are the most common malignant brain cancers in children, in whom brain tumors constitute the most common solid cancer [1]. Leptomeningeal dissemination of medulloblastoma, i.e., dissemination to the arachnoid, pia and cerebrospinal fluid (CSF), can occur in up to 40% of medulloblastoma patients at diagnosis and is found in most at recurrence [2-4]. Leptomeningeal medulloblastoma poses a dual challenge: 1) patients face poor prognosis despite intensive therapy, and 2) the small proportion of cured patients suffer serious long-term treatment-related sequelae, causing impaired quality of life and a serious burden to society, to their families and to themselves [1, 2, 5, 6]. Thus, leptomeningeal medulloblastoma requires development of more effective therapy.

The poor prognosis of leptomeningeal medulloblastoma is partially due to the challenge of delivering drugs effectively into the CSF [7]. These challenges include 1) the blood brain barrier, which prevents achievement of therapeutic CSF levels with systemic use of many drugs unless used at high doses that cause unacceptable systemic toxicity [7], and 2) direct intrathecal drug delivery via infrequent lumbar punctures that may provide only limited leptomeningeal exposure [8], especially in view of the rapid CSF turnover (6 h in humans, 2 h in mice), rapid drug clearance and uneven distribution in the CSF [9, 10]. Nevertheless, delivery of drugs directly into the CSF can be an attractive modality due to the greater therapeutic concentrations in CSF that can be achieved with significantly lower systemic exposure and fewer systemic side effects [7]. Thus, it is thought that improved delivery of drugs to the CSF will be beneficial.

A Phase I clinical trial found that continuous intrathecal infusion of topotecan, a topoisomerase I inhibitor, was well tolerated, suggesting that such an approach may help to circumvent some of the challenges in treatment of leptomeningeal disease [11]. A relevant question is whether such delivery is safe and effective and whether the preferred schedule is bolus or continuous. We therefore compared efficacy of topotecan delivered directly into the CSF as daily bolus injection with similarly-delivered topotecan as continuous infusion, using a mouse model of human leptomeningeal Group 3 medulloblastoma. Here we report that continuous and bolus IVT topotecan into mice with leptomeningeal medulloblastoma yielded similar survival advantage, similar improved control of brain leptomeningeal spread and mild advantage in control of spine leptomeningeal disease for the bolus treatment. We also find that both IVT topotecan delivery methods were associated with localized brain necrosis. We discuss possible limitations and approaches to improve the efficacy of topotecan delivery into the CSF.

## Materials and methods

### Cells

D425 Med medulloblastoma cells were a gift from Dr. Darrell D. Bigner (Duke University, Durham, NC) [12]. These cells were transduced with SMPU-R-MND lentiviral vector [13, 14] containing firefly luciferase and stable clones were selected by limiting dilutions and subsequent luciferase assay. D283 Med medulloblastoma cells stably expressing firefly luciferase in Luc(ff):zeocin/pcDNA3.1(+) (pJ00778) following selection in zeocin were a gift from Dr. Michael Jensen [15]. Both lines are classified as belonging to molecular subgroup 3 of medulloblastoma [16-20]. D425 were cultured in Ham’s F-12 medium containing 10% fetal bovine serum in a 37°C, 5% CO_2_ incubator. D283 were cultured in DMEM medium containing 10% fetal bovine serum and 0.6 mg/ml zeocin. Cell lines were negative for mycoplasma and were authenticated by small tandem repeats in November 2017.

Treatment of cultured cells with chemotherapy was performed as described in the legend to Figure 1. Bioluminescence was measured using a luminometer (Promega GloMax) after automatic injection of 100 μl of D-luciferin (0.33 mg/ml) into wells containing 100 μl of medium and cells.

### Reagents

Cytarabine, methotrexate and topotecan were purchased through the Children’s Hospital Los Angeles pharmacy. D-luciferin was from Biosynth International, Inc.

### Mice

Mice were housed at The Saban Research Institute of Children’s Hospital Los Angeles, a facility accredited by the Association for Assessment and Accreditation of Laboratory Animal Care International. All mouse procedures were approved by the Children’s Hospital Los Angeles Institutional Animal Care and Use Committee (protocol number 190) and were performed in strict accordance with recommendations of the latest (eighth) edition of the *Guide for the Care and Use of Laboratory Animals*.

Mice used were female J:NU mice (homozygous for the *Foxn1^nu^* mutation; The Jackson Laboratory). Mice in the intraventricular (IVT) treatment groups were cannulated by the vendor at age 8 weeks into the lateral ventricle according to the vendor’s standard coordinates. Mice in the bolus IVT treatment group were implanted with standard straight cannulas (PlasticsOne, 26 gauge, cat# C315GS-5/SPC), and mice in the IVT osmotic pump group received 28 gauge cat# 3280PM/SPC cannulas. Mice were shipped at age 9 weeks.

On the first day of the experiment D425-ff-luc medulloblastoma cells (2 × 10^5^ saline-washed cells in 2 μl per mouse) were injected into the cisterna magna of the mice while they were under ketamine/xylazine anesthesia. In mice receiving osmotic pumps, this injection was immediately followed by subcutaneous implantation of the drug-or saline-containing pumps, which were connected to the IVT cannulas via short catheter tubing. These catheters contained saline so as to delay the start of drug entry into the CSF until the day following implantation, a time when the other treatments were also scheduled to begin. Analgesia was provided by ketoprofen prior to cisterna magna injection and followed by ibuprofen in the drinking water after injection. Treatment was daily for bolus-treated mice for the duration of the experiment with IVT injections being given over 3 minutes each time, or continuously for mice with pumps for a minimum of 28 days. We used model 2004 Alzet osmotic pumps, which have a reservoir of 200 μl, a target pumping rate of 0.25 μl per hour and a pumping duration of at least 28 days. The lot of pumps used in this experiment was measured by the manufacturer to average 0.22 μl per hour.

Mice were observed daily by laboratory personnel and animal facility personnel, all of whom are trained to recognize symptoms requiring euthanasia. All efforts were made to alleviate potential animal discomfort. Euthanasia was performed when mice showed signs of tumor or illness such as head tilt or other neurological deficits, hydrocephalus, abnormal posture or movement, lethargy, rough coat, abnormal breathing, weight loss, or other signs of distress. These endpoints for euthanasia and the cranial localization of medulloblastoma tumors precluded their size from exceeding the currently recommended limits for tumor size in mice. Euthanasia was performed by isoflurane inhalation until mice were deeply anesthetized and their respiration ceases followed by perfusion with normal saline.

Bioluminescence imaging (Xenogen IVIS^®^ 100) of mice was performed twice weekly under isoflurane anesthesia after an intraperitoneal (IP) injection of D-luciferin (75 mg/kg body weight) as described [21]. Bioluminescence (radiance) is presented in the figures as photons/sec/cm^2^/steradian.

### Pathology

Mice were perfused with phosphate buffered saline and brains and spines were fixed in formalin overnight, paraffin-embedded, sectioned and stained with hematoxylin and eosin.

### Results

Medulloblastomas from molecular subgroup 3 are the ones most often found to have leptomeningeal spread [4, 22]. To choose human medulloblastoma cell lines for use in our leptomeningeal spread model we first tested chemosensitivity in culture of luciferase-expressing isolates of two medulloblastoma cell lines considered to belong to subgroup 3, D283 Med and D425 Med. We tested each line’s sensitivity to three chemotherapy drugs that can be used intrathecally: methotrexate, topotecan and cytarabine (ARA-C; Fig 1A-B). [23-28] Of the two cell lines both were resistant to methotrexate. D425 was only mildly sensitive to cytarabine (50%±6, SEM, cell kill at 250 μg/ml), and D283 showed resistance to it. For D425, incubation with 10 μg/ml topotecan for 3 days achieved 98%±0.2 cell kill, whereas D283 showed less than 50% cytotoxicity under those conditions. Upon comparing the sensitivity of three different clones of luciferase-expressing D425 to topotecan we found that all clones were similarly sensitive (Fig 1C) such that 0.1 μg/ml topotecan for four days induced 97-99% cell kill as measured by luciferase bioluminescence. We chose Clone #5 of D425 for the *in vivo* experiments.

**Fig 1.**
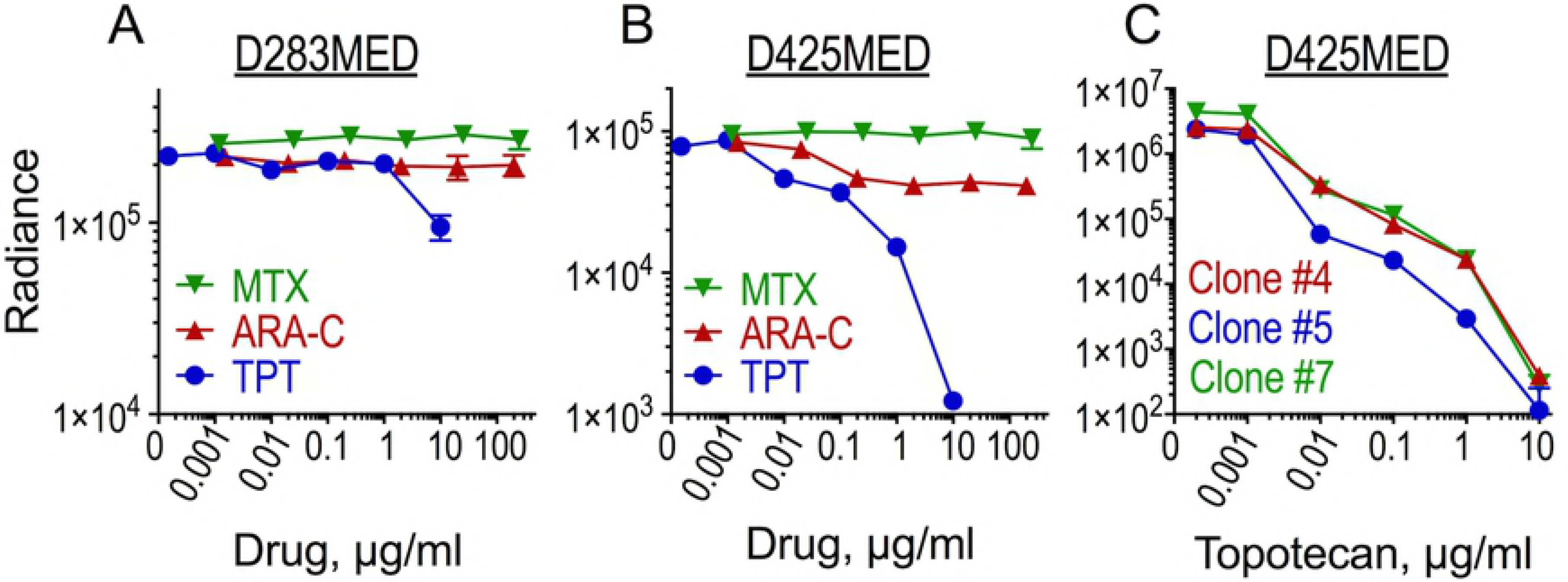
D425MED cells, but not D283MED, are sensitive to topotecan in culture. **(A-B)** D283 and clone 5 of D425 medulloblastoma cells, expressing firefly luciferase, were seeded at 2×10^3^ cells/well into 96-well plates, and methotrexate (MTX), cytarabine (ARA-C) or topotecan (TPT) were added for 72 h. Cells were analyzed for residual bioluminescence (Radiance) as a measurement of cells surviving following treatment. Data are the averages of duplicate measurements of duplicate wells ±SEM. **(C)** D425 clones 4, 5 and 7 expressing firefly luciferase were seeded at 1×10^4^ cells/well into a 96-well plate and exposed to the indicated concentrations of topotecan for 96 h between days 2 and 6 after plating with treatments on day 2 and 4 after plating. Bioluminescence was assessed 6 days after plating. Data are the average measurements of quadruplicate wells ±SEM.

D425 medulloblastoma cells expressing firefly luciferase were injected into the cisterna magna of mice under anesthesia. Pumps were implanted in the relevant IVT-cannulated mice immediately following injection. Treatment for all groups began the day following tumor and pump implantation. Treatment groups were 1) saline IP bolus, 2) saline IVT by continuous infusion via pump, 3) topotecan IP as bolus, 4) topotecan IVT continuously via pump, or 5) topotecan IVT by daily bolus injection. The topotecan daily dose delivered into the CSF via the pump IVT was 5.28 μg/mouse in 5.28 μl/day. The bolus IVT dose was 6 μg in 6 μl administered daily by manual injection. The IP dose was 15 μg/day [29]. Controls received saline in similar volumes for each route of administration.

Of the 35 mice in the experiment all but one developed leptomeningeal tumor, as determined by bioluminescence (Fig 2 and not shown), by symptoms related to tumor and as confirmed at necropsy. One mouse of the five in the IVT saline pump control group was still healthy appearing and gaining weight on day 46, three days after the last mouse in the whole experiment had been euthanized for tumor-related symptoms. On necropsy its brain showed no tumor, consistent with the absence of bioluminescence signal. Since all other saline control mice had extensive tumors and symptoms necessitating euthanasia between day 15-24, and even mice in the treatment groups all had obvious tumors by day 43, we concluded there was no tumor take in this mouse and excluded it from all figures and analyses.

**Fig 2.**
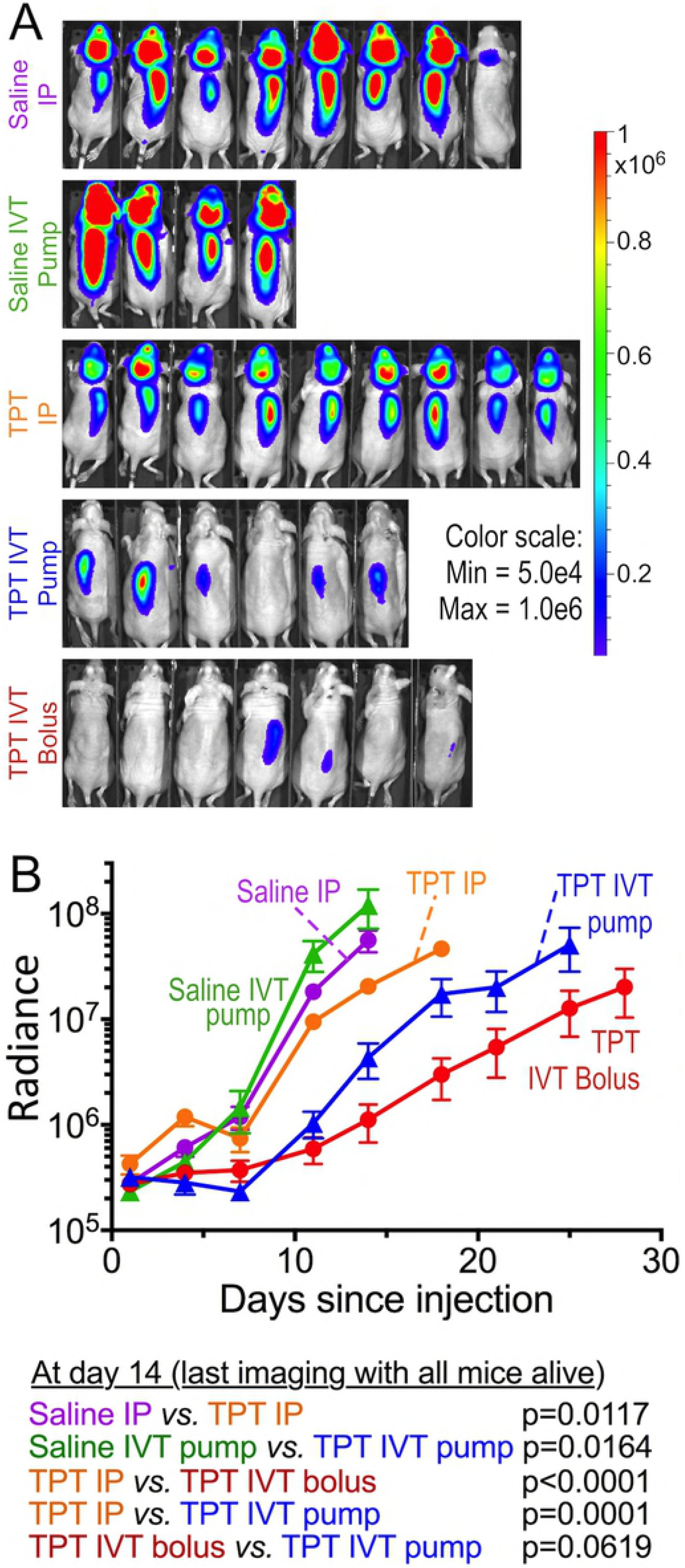
IVT topotecan slows leptomeningeal growth of D425 medulloblastoma cells in nude mice. D425-ff-luc cells were inoculated into the cisterna magna of nude mice. The following day treatment was started with topotecan via the indicated route. Bioluminescence was evaluated twice a week until mice showed clinically apparent signs of tumor on exam, at which time they were euthanized. (**A**) Bioluminescence imaging at day 14, which was the last imaging session when all mice in all groups were still alive. (**B**) Mean ± SEM of bioluminescence of each group. Means represent evaluations when all mice in the group were still alive, after which the curve is no longer shown. Below are p-values (log rank) comparing between the groups on day 14, which was the last imaging session when all mice in all groups were still alive. Saline IP, n=8 mice, Saline IVT pump, n=4, TPT IP, n=9, TPT IVT pump, n=6, TPT IVT bolus, n=7.

Mice in the saline control groups, whether via IP bolus injection or IVT via pump, fared worse than all topotecan groups in terms of having the most rapid increase in bioluminescence (Fig 2) and shortest survival (Fig 3). Among mice receiving topotecan, both groups receiving topotecan IVT showed slower rise in total tumor burden (measured by bioluminescence) and longer symptom-free survival compared to those receiving topotecan IP (Figs 2 and 3). Median survival was similar in mice receiving topotecan IVT by daily bolus compared to continuous delivery using the pump (Fig 3). The increase in total body bioluminescence of mice in the bolus compared to continuous (pump) IVT topotecan groups showed a trend towards slower rise in bioluminescence in the bolus group (Fig 2, day 14 p=0.0619, day 18 p=0.045, later p-values not significant).

**Fig 3.**
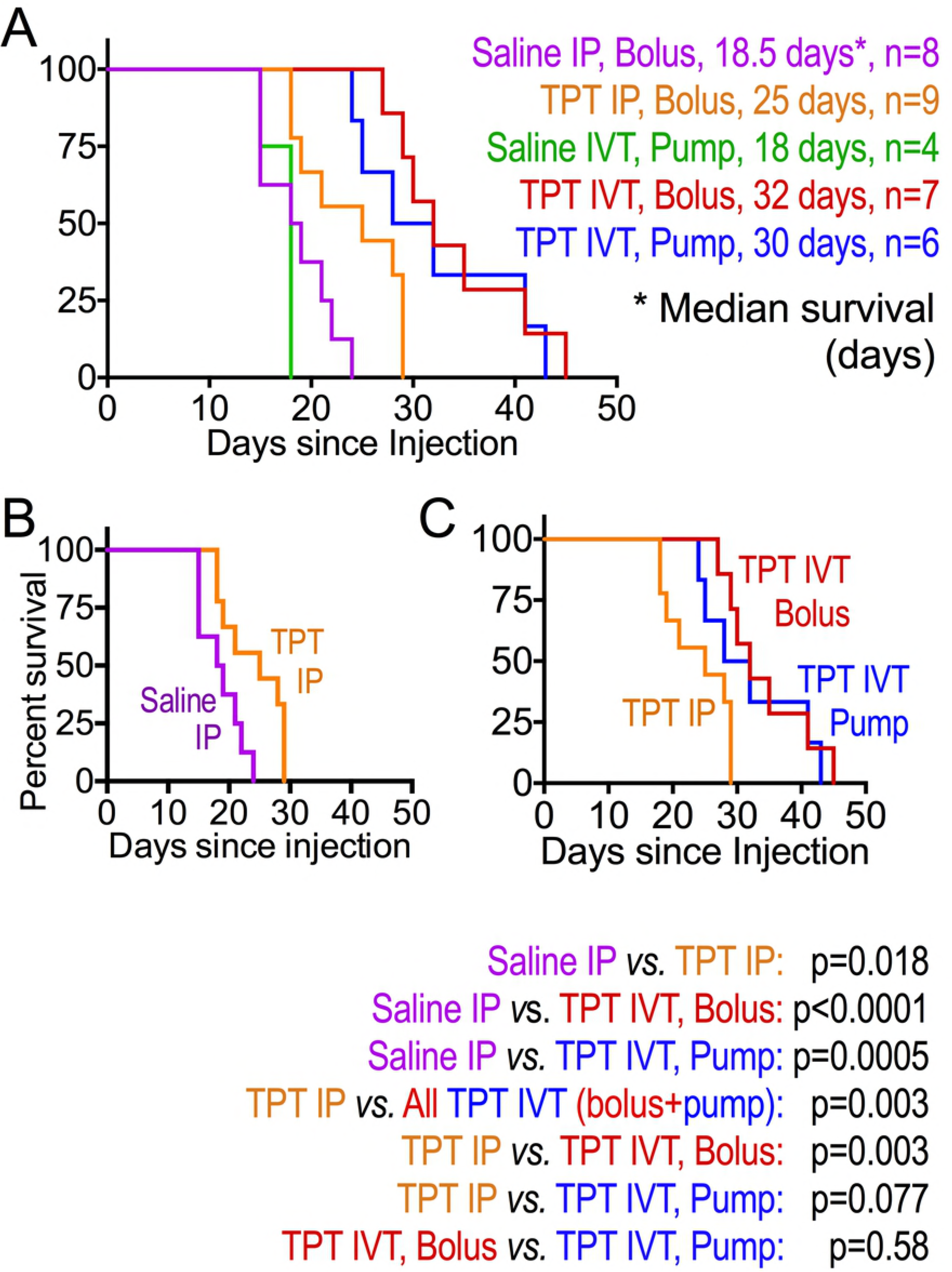
IVT topotecan delivered by daily injection or by continuous infusion similarly prolong survival of mice with leptomeningeal D425 medulloblastomas, prolonging survival compared to IP topotecan. Kaplan-Meier survival curves of mouse groups are shown. (**A**) comparison of all groups. (**B**) Daily IP topotecan *vs.* daily IP saline control. (**C**) Daily IP topotecan, daily IVT topotecan, or continuous IVT topotecan infusion via pump. Mice were euthanized when they showed clinical symptoms of tumor. Median survival of each group and its number of mice are noted to the right of panel (A). p-values, calculated by log rank, are shown below the survival panels.

We noticed that bioluminescence of the spines of mice receiving IVT topotecan rose faster than that of their brains, in which bioluminescence remained low (Fig 2A and not shown), suggesting that tumor in the spine was less responsive to IVT topotecan compared to the brains. This was different than mice treated with IP topotecan and the two saline groups, where tumor progression in each mouse was grossly similar in the spine and the brain. Plotting the ratio of spine to brain radiance confirmed that the increase in tumor load in brains of mice receiving topotecan IVT by either pump or bolus was indeed slower than in their spines, whereas in the other groups both rose similarly, as manifest in a steady ratio of spine-to-brain radiance (Fig 4A). Among the topotecan IVT-treated mice, radiance increase in the brain was slower in the bolus IVT group compared to the continuous infusion (pump) IVT group (Fig 4B). Spine tumor progression in mice receiving topotecan IVT by bolus showed a trend towards slower tumor growth compared to those receiving it by pump but did not reach statistical significance (Fig 4C). Thus, IVT topotecan was effective against leptomeningeal medulloblastoma in the brain itself, but less so in the spinal cord.

**Fig 4.**
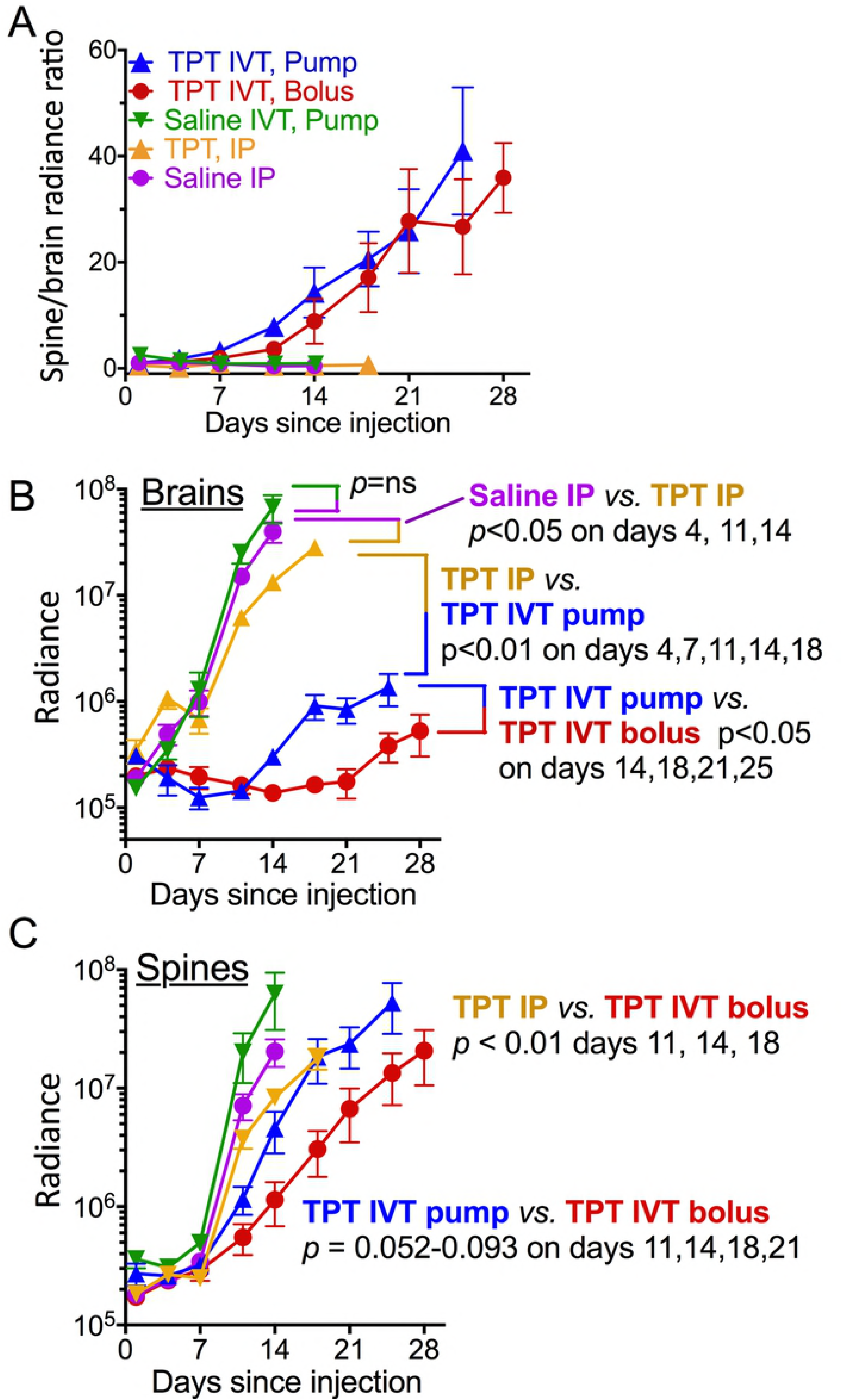
IVT topotecan preferentially slows leptomeningeal tumor growth in brains versus spines. Bioluminescence of brains and spines were calculated separately for each time point. Shown are mean ± SEM for each group, up to the date of first death in each group. **(A)** Ratios of spine-to-brain radiance measurements illustrate the relatively-faster increase in spine radiance compared to brain radiance in IVT TPT groups compared to the non-IVT groups. **(B)** Brain radiance measurements reveal more effective suppression of tumor growth in brains of TPT IP mice compared to saline IP in brains of TPT IVT (bolus or pump) mice compared to TPT IP and in brains of TPT IVT bolus mice compared to TPT IVT pump. **(C)** Spine radiance measurements reveal more effective tumor growth suppression in spines of TPT IVT bolus mice compared to TPT IP mice. There was a trend toward significance in spines of mice treated with TPT IVT bolus compared to TPT IVT pump, but it did not reach significance levels.

The hematoxylin and eosin (H&E)-stained sections of the brain and spinal column of control mice showed widespread diffuse leptomeningeal involvement of the cerebrum, cerebellum and spinal cord (Fig 5). There was extension of tumor cells focally into the Virchow Robin spaces of the brain and perineural involvement of cranial nerves and spinal nerve roots as well as surrounding dorsal root ganglia. The neoplastic cells were moderately pleomorphic and were characterized by markedly enlarged nuclei with prominent eosinophilic nucleoli and scant to moderate amounts of eosinophilic cytoplasm. The mitotic rate was brisk and there were frequent karyorrhectic cells.

**Fig 5:**
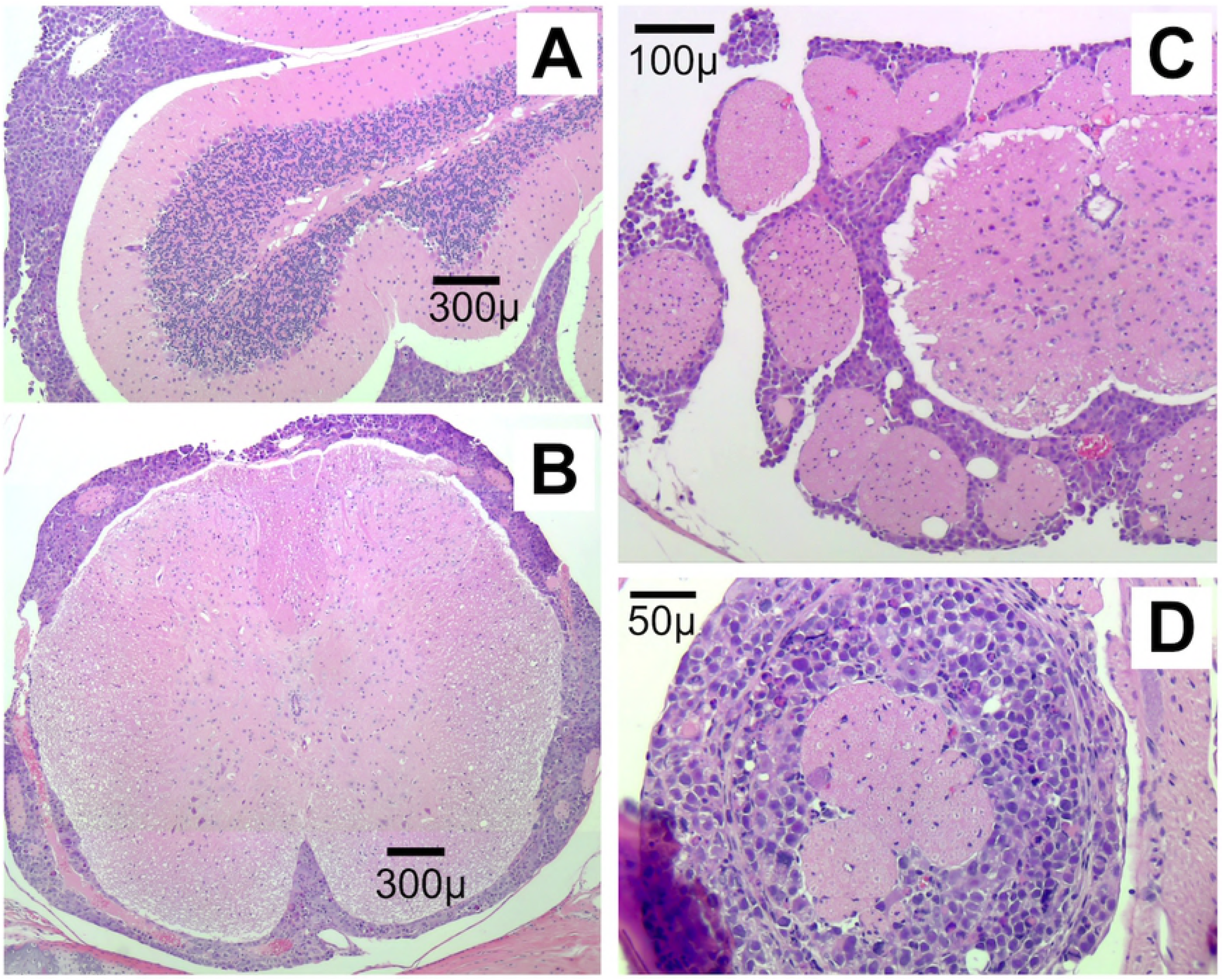
Leptomeningeal spread of D425 medulloblastoma cells is extensive. H&E stain of cerebellum (**A**; sagittal section) and spine (**B-D**; cross-sections) from a control mouse (IVT saline pump) euthanized at the time of tumor symptoms. Sections show extensive leptomeningeal spread of tumor cells around the brain and the spinal cord.

Consistent with the bioluminescence imaging, brains of mice receiving topotecan IVT showed very little tumor on H&E, although they had abundant tumor surrounding their spinal cords (Fig 4 and not shown). Mice receiving IVT topotecan showed varying degrees of inflammation and ventriculitis (Table 1).

**TABLE 1:** Brains of mice receiving IVT topotecan show inflammation and brain necrosis. Summary of findings in the harvested mouse brains, as evaluated by H&E staining of 2-4 FFPE sections from each brain: Numbers in the denominator reflect the number of brains from the group that were assessible for necrosis or inflammation. The numerator reflects the number of brains in which necrosis or inflammation was found. Evaluation was by two independent blinded observers.

|  | # of mice/<br>group | Necrosis | Inflammation |
| --- | --- | --- | --- |
| Saline IVT pump | 4 | 0 / 4 | 0 / 4 |
| Saline IP bolus | 8 | 0 / 8 | 0 / 8 |
| TPT IP bolus | 9 | 0 / 9 | 0 / 9 |
| TPT IVT bolus | 7 | **4 / 4<sup>a</sup>** | **5 / 7** |
| TPT IVT pump | 6 | **6 / 6** | **6 / 6** |

Mice receiving topotecan, regardless of route, did not demonstrate overt clinical systemic toxicity, as reflected in their normal behavior, typical feeding and comparable weight gain during the bulk of the experiment. Symptoms requiring euthanasia were those usually attributed to brain tumor-associated symptoms (weight loss, lack of grooming, hunched posture) but not symptoms one would anticipate with symptomatic spinal cord metastases such as paralysis or limb weakness. IVT topotecan mice had at least 1 log lower brain bioluminescence and less intracranial tumor in their brain sections compared to non-IVT topotecan mice (Fig 4B and not shown). It was therefore surprising that despite the lower tumor load within their brains (Fig 4B), these IVT topotecan mice showed only mild survival advantage (Fig 3A) and their euthanasia was prompted by brain-related symptoms. Histologic examination of brains of these IVT topotecan mice showed areas of brain necrosis in the cortical region above the hippocampus (Fig 6). No necrosis was seen in other brain regions nor in any of the control mice, including mice who received IVT saline via osmotic pump and those who received topotecan intraperitoneally. This indicated that the necrosis was not related to the presence of the cannula *per se* but, rather, to treatment with IVT topotecan. We cannot exclude that cannula termination position may have also played a role. Necrosis was more extensive and severe in mice treated IVT using the osmotic pumps compared to the IVT bolus-treated mice.

**Fig 6.**
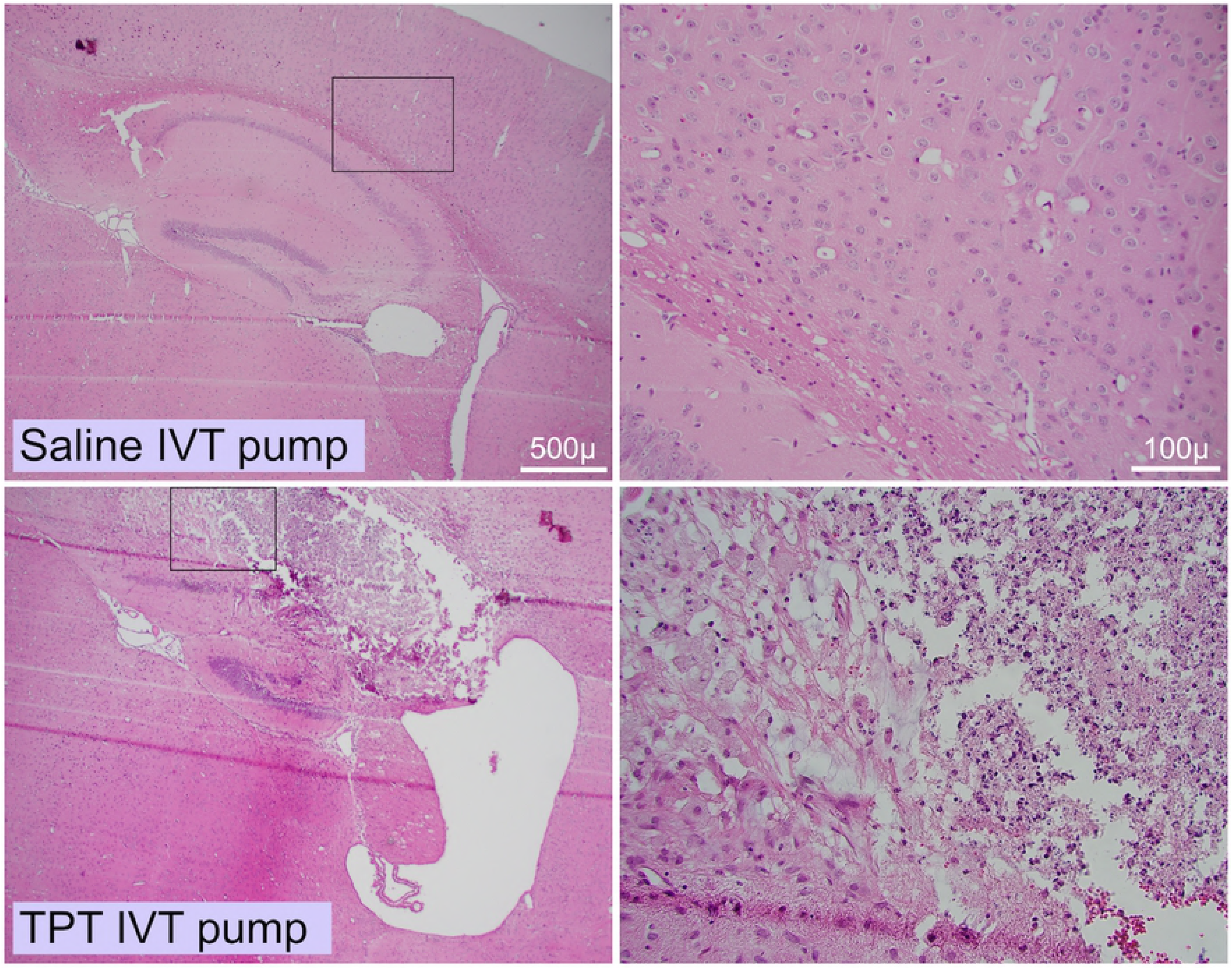
IVT topotecan delivered via intracranial cannula causes brain necrosis in a cortical region superior to the hippocampus. Representative H&E stained sections of brains from mice that received either IVT saline (top) or IVT topotecan (bottom) by continuous infusion pump. Brains of the other assessible mice that received IVT topotecan by bolus or by pump, but not of mice that received saline or IP topotecan, also showed necrosis in the region superior to the hippocampus. Magnifications are 40X (left) and 200X (right).

In summary, topotecan delivered into the cerebrospinal fluid prolonged symptom-free survival of mice in a leptomeningeal model of a Group 3 medulloblastoma using D425 Med cells compared to saline controls and to IP delivery of topotecan. Both IVT topotecan groups showed better tumor control within their brains compared to their spines with a trend toward better tumor control in brains of the bolus compared to the pump IVT topotecan mice. Under these conditions daily bolus IVT topotecan provided survival benefit that was similar to continuous IVT delivery and both were associated with varying degree of localized brain necrosis. The survival benefit of IVT topotecan may have been greater if the presumed locally-toxic effect of the topotecan could be averted.

### Discussion

The cultured medulloblastoma cell lines were variably sensitive to topotecan, a topoisomerase I inhibitor (Fig 1), similar to what others have reported [29-32]). Topotecan has clinical activity against childhood medulloblastoma in humans at concentrations above 1 ng/mL in CSF and exposure of over 8 h per day [29, 33]. Clinical trials have tested intrathecal bolus dosing of topotecan to determine its optimal dose, revealing limitations related to suboptimal drug level or toxicity at peak doses when using bolus dosing [7, 8, 34-36]. Continuous infusion of topotecan into the CSF is tolerable [11], but it is not yet known whether this method is more effective than bolus dosing. Here we report that in mice, topotecan showed only limited activity against leptomeningeal D425 Group 3 human medulloblastoma cells when delivered intraperitoneally. Topotecan produced a greater survival benefit when delivered directly into the CSF, either as continuous infusion using an osmotic pump or by bolus injection.

A prior study showed activity of IP topotecan against D425 subcutaneous xenografts when it was used at 1.9 mg/kg/day (47.5 μg per day for a 25 g mouse) 5 days/week x 2 weeks, a dose that was lethal to 10% of the mice [31]. We therefore based our dosing on a study to determine optimal curative dosing in a xenograft model of human ovarian cancer in nude mice, that produced no toxic deaths (0.625 mg/kg/day (15.6 μg per day for a 25 g mouse) 5 days/week x 4 weeks) [37]. In our study, IP topotecan daily at 15 μg per mouse per day slowed tumor growth (i.e., slowed the increase in bioluminescence) and prolonged median survival of mice carrying leptomeningeal D425 without overt clinical toxicity. The human equivalent dose is 1.27 mg/m^2^/day. This mouse dosing is in line with considerations extrapolated from pediatric topotecan dosing where a topotecan regimen of 1.2 mg/m^2^/day x 5 days systemically was well tolerated in children with neuroblastoma [38] and within the range considered tolerable and effective as studied in adults with ovarian cancer and small cell lung cancer [39, 40]. While IP topotecan prolonged median survival of our mice by 35% compared to IP saline (25 days *vs.* 18.5 days, respectively, p=0.0175; Fig 3), this approach did not achieve cures.

To achieve higher CSF topotecan and avoid systemic toxicity we tested direct intraventricular delivery into the cerebrospinal fluid by daily bolus and by continuous infusion. Dosing was based on published experience in pediatric patients and on our topotecan sensitivity experiments in D425 Med. In children, the maximal tolerated dose of bolus intrathecal topotecan is 0.4 mg/dose x 2 per week for 4 weeks [34]. A relatively well tolerated daily intrathecal bolus topotecan dose in children is 0.2 mg/day x 5 days [8]. Continuous infusion topotecan at that dose (0.2 mg/day x 7 days) was also well tolerated without signs of ventriculitis [11]. A 6-month-old Japanese infant was reported to receive 0.3 mg x 2 per week for 4 weeks followed by 0.4 mg x 1 per week for 1 month and then 0.4 mg less frequently for 12 additional months without severe arachnoiditis other than fever [28]. After calculating the volume of CSF in this 6-month-old infant to be approximately 16 ml, given an estimated weight of 8 kg [42] and a CSF volume of 2 ml per kg body weight [43], a 0.4 mg dose of topotecan in this patient would translate to a topotecan concentration in CSF of 25 μg/ml, which is somewhat higher than the concentration required to kill D425 cells in our cell culture experiments (1-10 μg/ml). In our continuously pumped IVT topotecan mice, we gave a 5.28 μg dose over a 24 h period, or 0.22 μg/h. Thus the maximum concentration of topotecan in the CSF (35 μl volume [41]) of pumped IVT mice after an hour of infusion might be calculated to be 6.29 μg/ml (0.22 μg / 35 μl), although the steady-state concentration will be lower due to CSF production and turnover (18 μl/h [41]). The similar daily dose of topotecan IVT (6 μg) delivered as bolus is expected to generate a short period with a very high concentration of drug in the CSF of mice in the bolus IVT group (171 μg/ml). The differences in maximum achieved drug concentrations between the two IVT topotecan groups may account in part for the better tumor control in the brains (Fig 4B) and the trend towards improved control in the spines (Fig 4C) of the bolus IVT topotecan mice compared to the pumped IVT group.

Since the pumped IVT dose (5.28 μg/day) would deliver higher drug amount to the CSF compared to the systemic (IP) topotecan (15 μg/day), it is not known if the higher efficacy of IVT topotecan was due to the route of drug delivery or the higher targeted dose of the IVT delivery. Since survival was similar in the IP saline control group and the IVT pump saline control group, this suggests that absence or presence of IVT catheter did not by itself affect survival. A minor limitation of the study is that the pumps, designed to reliably deliver drug for at least 28 days, were not replaced with new pumps after that time, since by then half the mice had to be euthanized due to tumor. As a result, it is formally possible that the three remaining mice in the pump group (euthanized on days 32, 41 and 43) had less drug delivered toward the end of the experiment.

We found that tumor was well suppressed within the brains of both the bolus and pump topotecan IVT groups, compared to the other groups, but less so in the spines. The IVT bolus delivery showed better control of the brain radiance and a trend toward better control of the spine radiance compared to continuous infusion of topotecan into the CSF. As mentioned above, a slightly higher daily dose of topotecan in the bolus group versus the pump group may have played a role in this, as could the higher peak dose of topotecan in the bolus group. The trend to lower control of the spine radiance in the continuous IVT topotecan group is also consistent with the thought that slow continuous drug infusion into the CSF may not achieve optimal CSF distribution due to the slow complex CSF flow through the heterogeneous CSF space [44, 45]. It suggests that a better distribution of drug to the spine may occur with the bolus injections. It is thus possible that a pulsatile and frequent intermittent flow that creates greater infusional forces may be more effective in increasing CSF mixing and optimizing drug distribution to the spine [44, 45] while also maintaining improved drug exposure over time. This confirms that topotecan can slow D425 Med xenograft growth in the brains of this leptomeningeal model using either continuous or bolus IVT modes of delivery, but that similar control of tumor growth in the spines will presumably require more effective delivery to this area.

The median survival times of the bolus and continuous IVT topotecan groups were similar, and both were significantly longer than the saline groups or the IP topotecan group. Since tumor burden within the brains of the IVT topotecan mice was much lower than in the other groups (Fig 2A, 4B) and we found localized necrosis in similar brain regions in both IVT topotecan groups (Fig 6, Table 1), we suspect that direct topotecan toxicity (e.g., brain necrosis) may have contributed to the demise of these mice. Despite the localized area of necrosis in the brain parenchyma, adjoining areas, including the ventricular lining, were unaffected. Relevant to this, convection enhanced delivery of topotecan into pig brain was reported to induce parenchymal damage in the brains as evidenced by magnetic resonance spectroscopy, with their histology showing necrosis along the catheter track [46]. The localized brain necrosis in our mice was only seen in those treated with IVT topotecan (i.e., those with both cannulas and topotecan) and was found in similar brain regions in them. While this suggests possible seeping of drug around the cannula as hypothesized in the pig brains above [46], we cannot rule out that cannulas which inadvertently terminated within the brain parenchyma may have contributed to the toxicity in some of the IVT topotecan mice.

In summary, we showed that prolonged delivery of topotecan directly into intraventricular CSF of mice with leptomeningeal D425 medulloblastoma effectively slows leptomeningeal tumor growth within the brain, is less effective in the spine, confers survival advantage on the mice, but is insufficient to cure them. The trend towards better control of the spine tumors in the bolus compared to the continuous IVT topotecan group suggests that pulsatile intermittent dosing into the CSF may improve drug distribution and anti-tumor effect.

## Acknowledgements

This work was supported by grant NS088965 from the NIH, National Institute of Neurological Disorders and Stroke and in part by the USC Coulter Translational Research Partnership Program. It was also supported in part by support from the Barbara Mandel Family Fund, the Brad Kaminsky Foundation Heroes of Hope Race, Grayson’s Gift and the Rachel Ann Hage Foundation.

## Abbreviations

CSF: cerebrospinal fluid
IP: intraperitoneal
IVT: intraventricular
TPT: topotecan

